# Lineage specific core-regulatory circuits determine gene essentiality in cancer cells

**DOI:** 10.1101/609552

**Authors:** Benedikt Rauscher, Luisa Henkel, Florian Heigwer, Michael Boutros

**Affiliations:** German Cancer Research Center (DKFZ), Division Signaling and Functional Genomics and Heidelberg University, Heidelberg, Germany

**Keywords:** cancer gene essentiality, transcriptional regulation, genetic networks, CRISPR, functional screens

## Abstract

Cancer cells rely on dysregulated gene expression programs to maintain their malignant phenotype. A cell’s transcriptional state is controlled by a small set of interconnected transcription factors that form its core-regulatory circuit (CRC). Previous work in pediatric cancers has shown, that disruption of the CRC by genetic alterations causes tumor cells to become highly dependent on its components creating new opportunities for therapeutic intervention. However, the role of CRCs and the mechanisms by which they are controlled remain largely unknown for most tumor types. Here, we developed a method that infers lineage dependency scores to systematically predict functional CRCs and associated biological processes from context-dependent essentiality data sets. Analysis of genome-scale CRISPR-Cas9 screens in 558 cancer cell lines showed that most tumor types specifically depend on a small number of transcription factors for proliferation. We found that these transcription factors compose the CRCs in these tumor types. Moreover, they are frequently altered in patient tumor samples indicating their oncogenic potential. Finally, we show that biological processes associated with each CRC are revealed by analyzing codependency between lineage-specific essential genes. Our results demonstrate that genetic addiction to lineage-specific core transcriptional mechanisms occurs across a broad range of tumor types. We exploit this phenomenon to systematically infer CRCs from lineage specific gene essentiality. Furthermore, our findings shed light on the selective genetic vulnerabilities that arise as the consequence of transcriptional dysregulation in different tumor types and show how the plasticity of regulatory circuits might influence drug resistance and metastatic potential.

## INTRODUCTION

Cancer cells rely on altered gene expression programmes to achieve optimal conditions for proliferation and survival and maintain their malignant phenotype (Califano and Alvarez, 2017). Previous studies suggested that small sets of transcription factors (TFs) can act together to form core-regulatory circuits (CRCs) that critically influence a cell’s transcriptomic state (Boyer et al., 2005). These TFs collectively control each other’s gene expression to form highly inter-connected autoregulatory loops (Odom et al., 2004, 2006; Saint-André et al., 2016). Changes in the regulation of CRC TFs can result in an altered gene expression state that can promote disease (Hnisz et al., 2013; Sanda et al., 2012). Therefore, it is important to identify CRCs and study their regulation to better understand disease mechanisms and find potential routes for novel therapeutic intervention.

While much progress has been made to define the CRCs for a number of cell types (Boyer et al., 2005; van Groningen et al., 2017; Saint-André et al., 2016; Sanda et al., 2012), the composition of CRCs across different cancer types and how they are regulated to promote tumorigenesis remains poorly understood. This is in part due to substantial experimental efforts that are required for the identification and validation of CRCs. Previous approaches have primarily employed ChIP-seq experiments to map the binding sites of TF genes in close proximity to super enhancer (SE) regions (Parker et al., 2013; Whyte et al., 2013). A recent study has shown that MYCN-amplified neuroblastoma cells crucially depend on the TFs that constitute their CRC for cell growth and survival (Durbin et al., 2018). Additionally, we and others have previously observed selective dependencies of other cancer cell types on TFs that play a role in development and differentiation of their associated tissues (Cheung et al., 2011; Kim et al., 2018; Rauscher et al., 2018). This indicated that CRC composition could critically influence selective gene dependencies in cancer cells of various tumor lineages.

Here, we examined whether we could systematically predict the CRCs of different cancer types from gene essentiality data. To this end, we analyzed genome-scale CRISPR fitness screens (Shalem et al., 2014; Wang et al., 2014) in 558 cancer cell lines (Meyers et al., 2017) for gene dependencies specific to a cancer’s tissue of origin (henceforth referred to as ‘lineage’). Among these, we found a strong enrichment for TF genes, many of which had previously been described to be part of the CRCs of certain cell types. Where available, public ChIP-seq and super enhancer data confirmed that these TF can regulate each other’s activity and are associated with super enhancer elements. We queried large-scale cancer genome databases (Cancer Genome Atlas Research Network et al., 2013; Forbes et al., 2017) and found that predicted CRC TFs are frequently affected by somatic alterations. Previous studies have demonstrated that functionally related genes (e.g. members of the same protein complex or pathway) show highly correlated essentiality profiles across cell lines (Pan et al., 2018). We employed this strategy to group all lineage dependency genes into functional units and link them to predicted CRCs. This revealed biological processes implicated in the aberrant regulation of CRCs in cancer that could represent potential points of vantage for therapeutic interventions. Finally, we explored how invading tumour cells might be able to adapt their CRCs in order to adapt to their new niche. Our analysis shows that addiction to lineage-specific core-regulatory circuits and their dysregulated state occurs across a broad range of tumor types. It provides a foundation for studying these CRCs and how they control different tumor lineages and demonstrates how CRC composition determines tissue-specific gene and pathway dependencies.

## RESULTS

### Context-dependent essential genes are enriched for lineage dependency transcription factors

Context-specific essential genes are genes whose knockout has little or no effect on the proliferation and survival of most cell lines, while a small number of cell lines are highly dependent on their activity. We assumed that core-regulatory circuits are composed of transcription factors that are selectively essential in the associated tumour lineages. Therefore, we first tested whether TFs were overrepresented among all context-dependent essential genes. To this end, we identified all genes for which increased dependency - as measured in CRISPR fitness screens (Meyers et al., 2017) - was observed in a restricted set of all cancer cell lines (Figure 1A, Figure 1B, Supplementary Figure 1A). We then ranked all genes by the number of selectively dependent lines. The oncogenes *KRAS, BRAF, NRAS* and *PIK3CA* were among the top ranking context essential genes (Figure 1C). Cells with mutations in these genes are well known to become selectively dependent on them through oncogene addiction indicating that we correctly identified context-dependent essentiality (Weinstein and Joe, 2008). Gene set enrichment analysis (Sergushichev, 2016) further revealed that TFs (Gene Ontology Consortium, 2015) were indeed overrepresented among the most context-dependent essential genes (NES = 1.45, FDR = 0.0017; Figure 1D).

**Figure 1.**
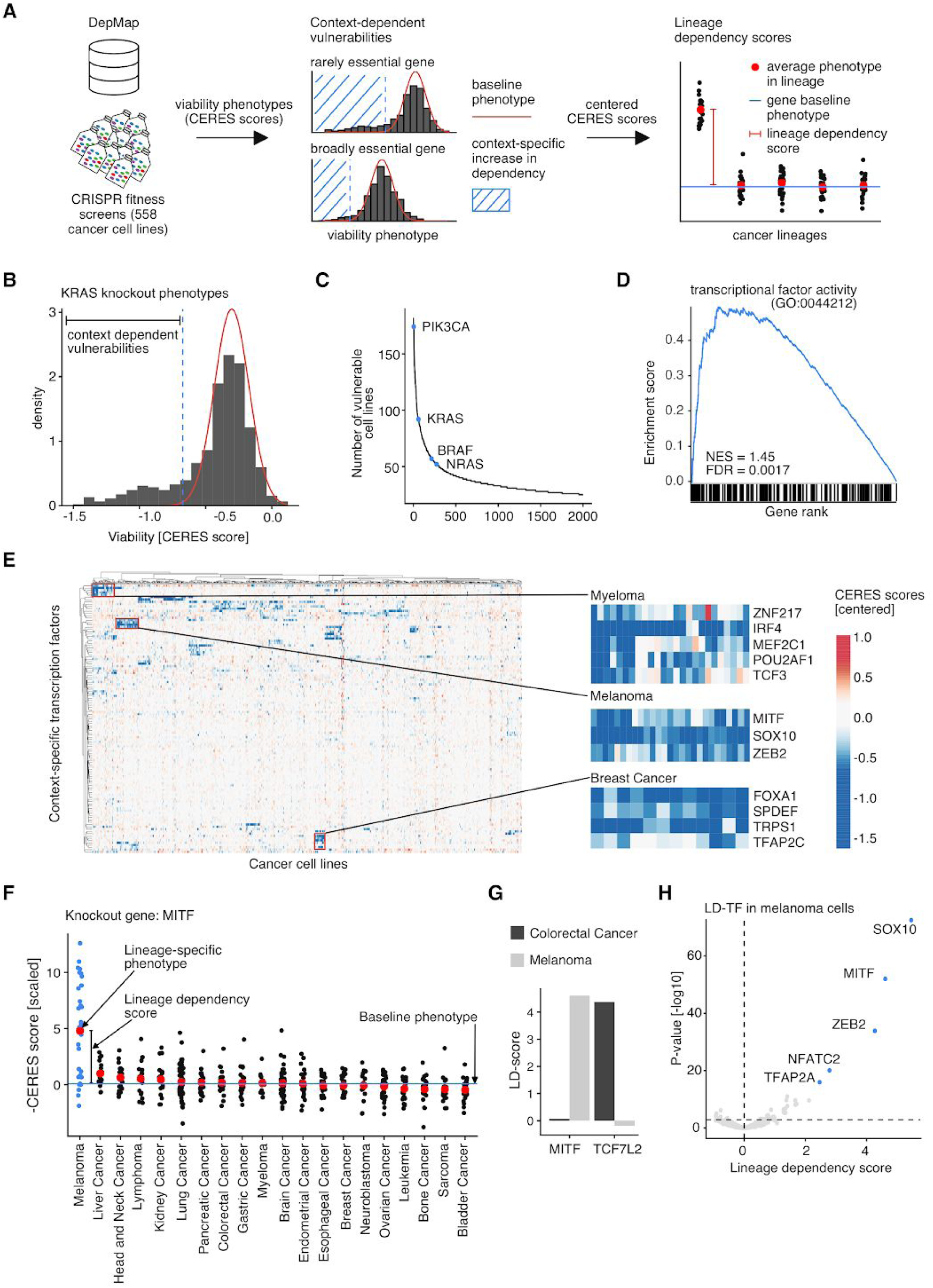
Context-dependent essential genes are enriched for lineage dependency transcription factors (LD-TF). (A) A schematic workflow illustrates the processing of CRISPR fitness screen phenotypes (CERES-scores; negative values indicate loss of viability) to determine context-specific vulnerabilities (middle) and lineage dependency genes (right). Context-specific phenotypes significantly deviate from the baseline distribution of knockout phenotypes for a gene across all cell lines. Genes where context-specific vulnerabilities are observed specifically in cell lines derived from a certain cancer lineage are termed lineage dependency genes. (B) A histogram of CERES scores measured upon KRAS knockout illustrates context-specific gene dependency in the case of oncogene addiction. The red curve indicates the estimated baseline phenotype. The vertical blue line marks a 20% FDR cutoff separating expected from context-specific phenotypes. (C) Genes are ranked by the number of cell lines in which they are classified as context-essential. Several well-characterized context-essential oncogenes are highlighted. (D) Genes that show a high degree of context-essentiality are enriched for transcription factors (TF; Gene Ontology Molecular Function). Genes are ranked by the number of cell lines that specifically depend on them. Black bars at the bottom highlight genes with annotated transcriptional activator activity (NES = normalized enrichment score; FDR = false discovery rate). (E) Clustering cell lines by context-essential transcription factors (n=117) reveals highly tumor type specific subgroups. Selected TF subgroups with specific phenotypes in myeloma, melanoma and breast cancer are highlighted on the right. (F) A dot plot visualizes cancer type specific fitness phenotypes upon knockout of the MITF transcription factor. Each dot represents a cell line (n=558). Red dots indicate the average phenotype for each cancer type and the horizontal line illustrates the median knockout phenotype across all cell lines. We call the distance between each group average and the blue median line the ‘Lineage dependency score’ (LD-score). (G) LD-scores for two selected example genes *MITF* and *TCF7L2* in melanoma (n=34) and colorectal cancer cells (n=27). (H) Volcano plot highlighting transcription factors with the strongest LD-scores in melanoma cells (n=34). Selected genes are highlighted in blue.

We next clustered cancer cell lines by their dependency on context essential TFs (Figure 1E). We found several distinct clusters of genes that showed highly specific essentiality in cell lines derived from the same tissue of origin. These findings prompted us to devise a score to quantify selective gene dependency of cancer lineages that we termed ‘Lineage dependency (LD)-score’. This score quantifies lineage dependency as the difference between the average viability phenotype upon gene knockout in each individual cancer lineage compared to the overall phenotype across all cancer types (Figure 1F). Next, we calculated LD-scores for all TF (Lambert et al., 2018) and non-TF genes in cancer lineages sufficiently represented in the CRISPR data set (n_cell lines / lineage_ > 10; Supplementary Figures 1B-D, Supplementary Table 1). To assess whether LD-scores could recapitulate known lineage-specific gene dependencies, we examined TFs with high melanoma LD-scores. As expected, we found the melanocyte master regulator *MITF (Garraway et al., 2005; Levy et al., 2006)* and the Wnt-signaling effector *TCF7L2* (Korinek et al., 1997; Morin et al., 1997; Zhan et al., 2017) among the top scoring genes for melanoma and colorectal cancer, respectively (LD-score = 4.61, FDR = 5.40×10^−49^ and LD-score = 4.37, FDR = 6.18×10^−31^; Figure 1G). Along with MITF the LD-score further identified other melanocyte-specific TFs known to bind the *MITF* promoter to regulate its gene expression, such as *SOX10* (LD-score = 5.46, FDR = 3.50×10^−69^), *PAX3* (LD-score = 1.71, FDR = 9.82×10^−7^) or *ZEB2* (LD-score = 4.27, FDR = 3.22×10^−31^) (Hartman and Czyz, 2015). These findings suggest that context-dependent essentiality can in fact reveal transcriptional master regulators of different tumor lineages.

### LD-TFs compose cancer lineage specific core-regulatory circuits

We next investigated whether LD-TFs represent, at least in part, the CRCs of the dependent cancer lineages. Thus, we selected significant LD-TFs (LD-score > 1, FDR < 5%; 205 in total) as components of putative regulatory circuits in the associated cancer lineages (Supplementary Table 1). The number of LD-TFs identified for each cancer lineage was highly variable ranging from 5 (lung cancer) to 60 (myeloma; Supplementary Figure 2A). Clustering of cancer lineages by LD-scores revealed groups of common developmental background (Supplementary Figure 2B). TFs acting together as CRCs are known to be highly inter-regulated (Boyer et al., 2005) and under transcriptional control of super enhancer (SE) elements (Hnisz et al., 2013). Therefore, we further annotated our putative CRCs with public TF-target-gene (Harmonizome) and super enhancer data (SEdb; Figure 2A) (Jiang et al., 2019; Rouillard et al., 2016).

**Figure 2.**
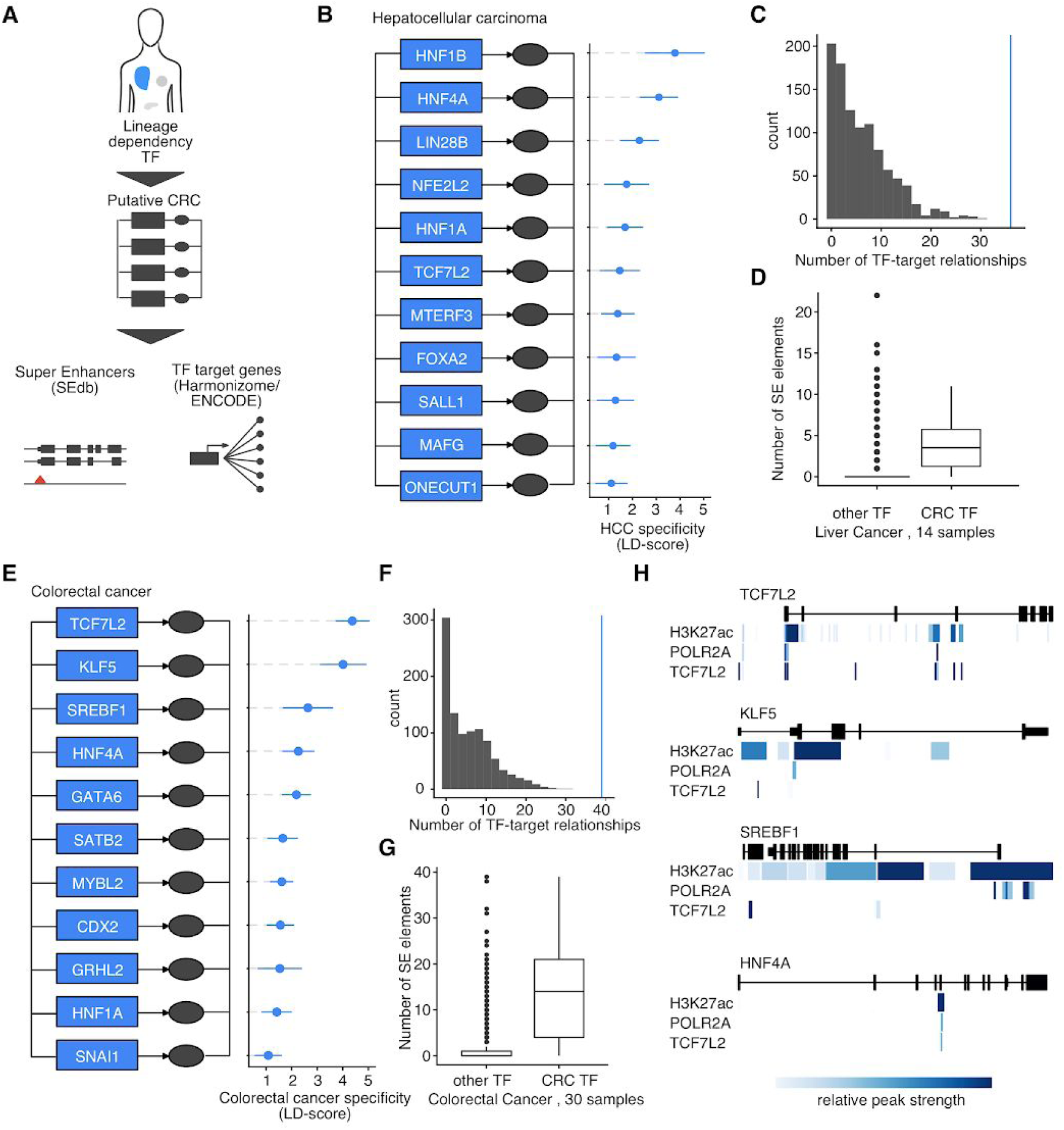
LD-TF compose cancer type specific core-regulatory circuits. (A) Lineage dependency transcription factors (LD-TFs) inferred from fitness screens were examined for regulatory relationships (TF targets as reported by Harmonizome/ENCODE) and associated super enhancer elements (as reported by SEdb) to predict cancer type-specific core-regulatory circuits. (B) The LD-TFs of hepatocellular carcinoma are part of the hepatocyte CRC. Blue boxes represent genes and ovals represent their protein products. Genes are arranged as interconnected circuits indicating regulatory relationships. The lineage dependency score (LD-score) is shown for each CRC gene. Error bars are ± 2 s.e.m. (C) A histogram of the number of TF target gene relationships reported in Harmonizome for 1,000 random samples of 11 transcription factors. The vertical blue line indicates the number of reported TF target gene relationships for members of the HCC CRC. (D) Number of super enhancer elements reported in SEdb for members of the HCC CRC compared to all other TF in liver tissue samples. The center of each box is the sample median, the whiskers extend from the upper (lower) hinge to the largest (smallest) data point no further than 1.5 times the interquantile range from the upper (lower) hinge. (E-G) The predicted CRC of colorectal cancer consists of 11 TF (LD-score > 1; FDR < 5%) and exhibits similar properties to the HCC CRC on TF-target gene and super enhancer data. (H) Public ChIP-seq data on H3K27ac, POLR2A and TCF7L2 target proteins shows POLR2A and TCF7L2 binding in enhancer regions that are found close to the transcription start sites of colorectal cancer CRC genes. Blue colour intensity represents the relative peak strength.

To validate this approach, we examined whether it could recapitulate known biology. The CRC of hepatocytes is one of the most well-studied regulatory circuits (Odom et al., 2004, 2006). Here, the hepatocyte nuclear factors *HNF1A, HNF4A* and *ONECUT1* (*HNF6*) as well as *FOXA2* have been described as key players regulating each other’s activity to maintain the hepatocyte transcriptional state. These TFs were also part of the hepatocellular carcinoma CRC we predicted for hepatocellular carcinoma from gene essentiality data (Figure 2B). In addition, our predicted CRC contained additional TFs that have previously been described (El-Khairi and Vallier, 2016; Nguyen et al., 2014) to play important roles during liver development, such as *HNF1B* (LD-score = 3.79, FDR = 8.66×10^−7^) or *LIN28B* (LD-score = 2.29, FDR = 9.06×10^−6^). As expected, public transcription factor databases (ENCODE Project Consortium, 2004; Lachmann et al., 2010; Matys et al., 2003; Rouillard et al., 2016; Xu et al., 2013) report a large number of TF-target gene relationships (i.e. genes can bind each other’s promoters) between TFs in our predicted liver CRC (p = 6.03×10^−7^, Kolmogorov-Smirnov test; Figure 2C). Also, the SEdb database (Jiang et al., 2019) reports a higher number of SE elements associated with these TFs compared to other transcription factors in liver samples (p = 1.73×10^−5^, two-sided Wilcoxon rank sum test; Figure 2D). This implies that LD-TFs could in fact recapitulate lineage specific CRCs.

We next examined the putative CRC of colorectal cancer that, as such, had not previously been described. In total we found 11 TFs with significant LD-scores in colorectal cancer cell lines (Figure 2E). These included several genes known to play a role during the development and differentiation of colon cells (Gao et al., 2009; Silberg et al., 2000; Yu et al., 2017) such as *CDX2* (LD-score = 1.56, FDR = 2.74×10^−6^) and *SATB2* (LD-score = 1.65, FDR = 1.12×10^−5^). Some of these genes have also been shown to promote tumorigenesis when aberrantly activated; *TCF7L2*, the gene with the highest colorectal cancer LD-score (LD-score = 4.37, FDR = 6.18×10^−31^) is the effector of the Wnt signaling pathway that is mutated in the majority of colorectal cancers. Again we found that TF-target gene relationships between colorectal LD-TFs had been reported frequently in the Harmonizome database (p = 1.09×10^−7^, Kolmogorov-Smirnov test; Figure 2F). Similarly, more SE elements linked to these TFs had been identified compared to other transcription factors in colon or colorectal cancer samples (p = 6.26×10^−7^, two-sided Wilcoxon rank sum test; Figure 2G). Moreover, public ChIP-seq data (ENCODE Project Consortium, 2004) suggest that TCF7L2 can recruit POLR2A to bind enhancer regions (H3K27ac bound DNA) proximal to the TSS of other colorectal CRC TF to promote gene expression in the colorectal cancer cell line HCT116 (Figure 2G, Supplementary Figure 2C).

### Lineage dependency transcription factors are important in both, normal and cancer tissue and can have oncogenic potential

Since CRCs inferred from LD-scores included several transcription factors that had previously been implicated in tumorigenesis and cancer progression, we asked whether LD-TFs could act as lineage survival oncogenes (Garraway and Sellers, 2006). Lineage survival oncogenes are genes that play crucial roles during tissue development and that can be deregulated in cancer (Garraway and Sellers, 2006). They are thought to i) play important roles in normal lineage survival during development, ii) be deregulated in cancers of the associated lineage and iii) be affected by somatic alterations in tumor subsets. To answer these questions for significant LD-TFs (LD-score > 1, FDR < 5%), we surveyed the genome databases GTEx, ExAC, TCGA and COSMIC (Figure 3A) (Cancer Genome Atlas Research Network et al., 2013; Forbes et al., 2017; GTEx Consortium et al., 2017; Lek et al., 2016). We found that damaging mutations in LD-TFs occur less frequently in living individuals compared to other transcription factors or non-TF genes (ExAC; p = 6.303×10^−10^, two-sample Student’s t-test; Figure 3B, Supplementary Figure 3A) (Lek et al., 2016) indicating that LD-TFs are subject to strong selective pressure. Further, gene expression of LD-TF genes was restricted to their associated tissues in non-cancer samples (GTEx; p = 2.2×10^−16^, two-sample Wilcoxon rank sum test, Figure 3C, Supplementary Figure 3B) (GTEx Consortium et al., 2017). Similarly we found higher expression of LD-TF genes in tumor transcriptomes representing the associated cancer lineage as compared to other lineages (TCGA; p < 2.2×10^−16^; two-sample Wilcoxon rank sum test, Figure 3D, Supplementary Figure 3C) (Cancer Genome Atlas Research Network et al., 2013). We tested whether LD-TFs were differentially expressed in cancer compared to healthy samples and found that LD-TFs were down-regulated less frequently (p = 0.11) and up-regulated more frequently (p = 0.10) than expected by chance (Figure 3E). For the remaining genes, however, we did not detect a change in expression. Also, many TFs that were differentially expressed between healthy and cancer samples did not show a phenotype in CRISPR screens (Supplementary Figure 3D-E) (Barretina et al., 2012; Meyers et al., 2017). Finally, we investigated whether LD-TFs are affected by somatic molecular alterations in tumors (Figure 3F) (Forbes et al., 2017). While we did not find frequent (>5% of samples corresponding to a tumor lineage) somatic variants for all LD-TFs, we identified common somatic alterations for a significant subset of LD-TFs (p = 2.235×10^−11^, Fisher’s exact test). These results indicate that a subset of LD-TFs could in fact act as lineage survival oncogenes, modify CRC activity and initiate a cancer promoting gene expression program.

**Figure 3.**
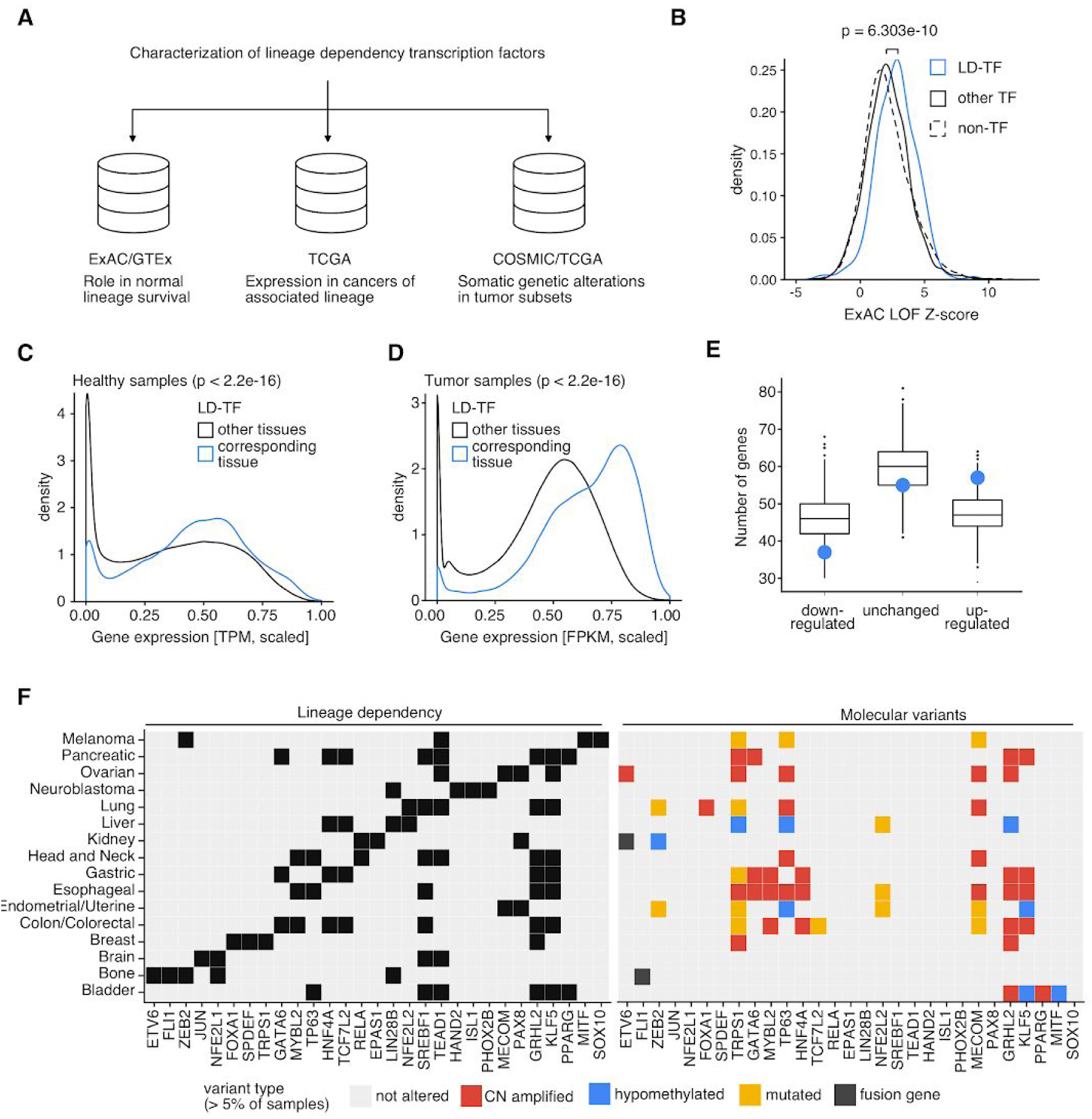
Lineage dependency transcription factors are important in both normal and cancer tissue and can have oncogenic potential. (A) Population and cancer genome databases were interrogated to characterize gene expression and molecular variants of LD-TFs in healthy and cancer tissues. (B) Damaging mutations in LD-TFs (blue curve; n=216) are found less frequently than expected compared to other transcription factors (solid black curve; n=1,182) and non-transcription factor genes (dashed black curve; n=15,010) according to the ExAC LOF score (two-sided Student’s t-test). (C) LD-TFs are highly expressed in their associated tissues (blue curve; n=86,140) compared to other tissues (black curve; n=1,398,510) in healthy tissue samples included in the GTEx database (two-sided Wilcoxon rank sum test). (D) LD-TFs are highly expressed in their associated tumor types (blue curve; n=71,411) compared to other tumors (black curve; n=1,186,919) in samples included in the Cancer Genome Atlas (TCGA; two-sided Wilcoxon rank sum test). (E) LD-TFs (blue dot) are down-regulated less often and up-regulated more often than expected (black box; 1,000 TF random samples) in tumor samples compared to healthy reference samples (TCGA). Differential gene expression was determined using a two-sample Welch’s t-test. The Benjamini-Hochberg method was used to control the false discovery rate at 20%. The center of each box is the sample median, the whiskers extend from the upper (lower) hinge to the largest (smallest) data point no further than 1.5 times the interquantile range from the upper (lower) hinge. (F) Molecular variants including somatic mutations, somatic copy number gains, hypomethylation or gene fusion occur more often than expected in LD-TFs (p = 2.627 × 10^−11^, Fisher’s exact test). A gene was determined as altered in tumor type if the alteration was found in more than 5% in the COSMIC database. The heatmaps show LD-TF status (left heatmap; black indicates lineage dependency at LD-score > 1 and FDR < 5%) and molecular alteration status (right heatmap) for the top 3 LD-TF determined for each cancer type that could be mapped to the COSMIC database.

### Co-dependency predicts mechanisms leading to CRC deregulation in cancer

Samples from tumors and healthy adjacent tissue represent two distinct transcriptomic states when clustered by principal component analysis (PCA) (Califano and Alvarez, 2017). A healthy cell has to adapt its gene expression program in order to transform into a cancer cell with increased cell proliferation and survival (Figure 4A). This can be achieved through the deregulation of a TFs with oncogenic potential (Bradner et al., 2017). Since oncogenic TFs are part of CRCs that control a cell’s transcriptomic state, we assumed a model where the aberrant activation of a TF with transformative potential leads to CRC deregulation; either through somatic alterations in an upstream pathway or the TF itself (Alvarez et al., 2016). This results in an altered transcriptional output that enables cancer development (Figure 4B). We expected highly selective dependency on components of the CRC and upstream pathways (Supplementary Figure 4A) required to achieve its deregulation, while we assumed that cancer promoting target genes should be less lineage specific. Again, we took melanoma cells as an example to explore this hypothesis since the mechanisms of aberrant *MITF* activation have been carefully studied in the past (Garraway et al., 2005; Levy et al., 2006; Wellbrock and Marais, 2005; Wellbrock et al., 2008). We first selected all genes (both TF and non-TF) with significant LD-scores in melanoma cell lines (LD-score > 1, FDR < 5%). In total, we selected 15 TFs representing the CRC and 161 non-TF genes as putative CRC regulators and target genes. We then clustered these genes by codependency across all 558 screened cell lines to group them into functional units as described previously (Figure 4C) (Pan et al., 2018; Rauscher et al., 2018). This revealed distinct clusters including a central cluster showing high codependency with a large fraction of selected genes. This cluster consisted of key regulators of the MAPK signaling pathway, such as *BRAF* (LD-score = 5.69, FDR = 3.76×10^−45^), *MAPK1* (ERK; LD-score = 3.16, FDR = 2.58×10^−28^) or *MAP2K1* (MEK; LD-score = 1.32, FDR = 6.55×10^−6^). It further included the LD-TFs *MITF* and *SOX10. BRAF* is mutated in the majority of melanomas and is well known to regulate *MITF* to promote tumor proliferation and survival (Wellbrock et al., 2008). In addition, we found clusters that were comprised of genes involved in processes such as epithelial-mesenchymal transition (EMT) or apoptosis (Figure 3C) (Cicchini et al., 2008; Mott et al., 2007). Both of these processes are known to be dysregulated as the consequence of aberrant *MITF* activation (Figure 4D) (Denecker et al., 2014; Hartman and Czyz, 2015; Levy et al., 2006). Encouraged by these observations we aimed to systematically link the CRCs of different tumor lineages to associated biological processes. We selected significant LD-genes for each cancer lineage and grouped them into 2-10 distinct codependency clusters (see Methods). We then performed a Gene Ontology enrichment analysis for each cluster (Alexa and Rahnenfuhrer, 2010; Gene Ontology Consortium, 2015; Ritchie et al., 2015). In total we identified 149 clusters across 20 cancer lineages that we linked to a diverse range of biological processes (1,508 unique GO IDs, FDR < 20%; Supplementary Figure 4B, Supplementary Table S2). The 7 clusters selected for melanoma, for example, enriched for processes such as MAPK signaling (p=1.40×10e^−4^), melanocyte differentiation (p=1.76×10^−4^), cell division (p=1.77×10^−3^), or apoptosis (p=3.32×10^−4^). As expected, specificity of dependency phenotypes (quantified by average LD-scores of genes in a cluster) was highest for CRC genes and their upstream regulators (MAPK signaling) while target processes showed weaker, although still significant, specificity (Figure 4E). These observations suggest that regulatory mechanisms of lineage specific CRCs can in fact be identified by examining global codependency profiles of lineage dependency genes.

**Figure 4.**
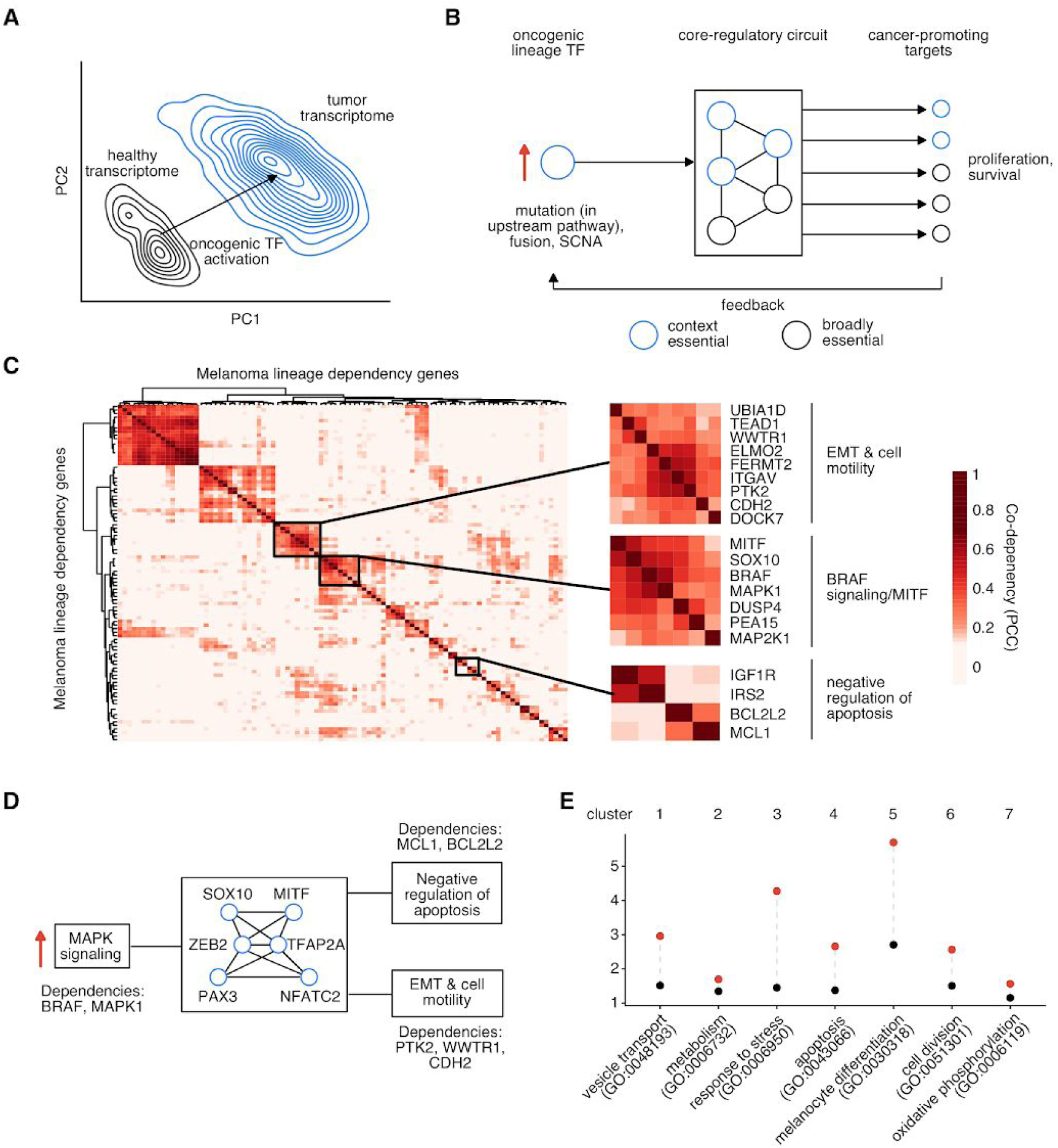
Lineage specific core-regulatory circuits (CRC) control the cell’s transcriptional state and determine context-specific gene essentiality. (A) A principal component analysis (PCA) plot depicts clustering of healthy (black lines) and tumor (blue lines) transcriptomes in patient colon/rectum samples (TCGA). Thi hints at a mechanism where a healthy cell’s transcriptome is transformed into a tumor transcriptome through the aberrant activation of an LD-TF with oncogenic potential. (B) In this model, one or few oncogenic LD-TFs control an interconnected core-regulatory circuit (CRC) consisting of a small number of other lineage dependency and broadly essential transcription factors. The CRC expresses cancer promoting target genes that can feed back into pathways upstream of the lineage dependency oncogene TF. The red arrow indicates an oncogenic up-regulation. (C) A codependency heatmap of can group melanoma lineage dependency genes into functional units revealing key regulatory mechanisms linked to the melanoma CRC. Clusters of genes associated with processes known to be regulated by the melanoma master regulator MITF are highlighted (EMT = epithelial-mesenchymal transition). (D) A schematic illustration shows part of the regulatory architecture of the melanoma CRC (center, TFs with high LD-scores are highlighted) that is revealed by codependency analysis. Selected gene vulnerabilities are highlighted. (E) A dot plot shows average (black dot) and maximum (red dot) LD-scores for genes associated with one of 7 codependency clusters of melanoma lineage dependency genes. A representative enriched GO Molecular Function term is shown for each cluster.

### Metastatic cancer cells might alter their core-regulatory circuit to adapt to their new niche

We noticed that in many cases dependency on LD-TF could vary across cell lines representing the same cancer lineage. For example, several melanoma cell lines do not depend on the melanocyte master regulator *MITF* for proliferation and survival. At the same time, *MITF* expression is decreased in these cell lines (Figure 5A). Similarly, MITF resistant melanoma cell lines show decreased expression of key melanocyte differentiation markers such as *MLANA, PMEL* or *TYR* (Figure 5B) (Vazquez et al., 2013). This phenomenon, termed ‘phenotype switching’, is not restricted to cell lines but is also observed in patient tumors (Supplementary Figure 5A) where it is associated with increased resistance to a range of therapies (Müller et al., 2014; Perotti et al., 2016) and increased invasive potential (Vachtenheim, 2017). We hypothesized that melanoma cells might have the ability to adapt their CRC in order to achieve a gene expression state that is more favorable for invasion. To investigate this, we clustered 5,471 tumor samples (Cancer Genome Atlas Research Network et al., 2013) by their expression of significant LD-TF genes (LD-score > 1, FDR < 5%) using t-Distributed Stochastic Neighbor Embedding (t-SNE). The expression of LD-TF genes was sufficient to achieve clustering of tumor samples by cancer type (Figure 5C, Supplementary Figure 5B). We noticed that, while all MITF_high_ samples clustered together to form a melanoma-specific cluster (Figure 5C), many MITF_low_ samples clustered with soft tissue sarcoma samples instead. Loss of MITF expression was associated with increased expression of the key EMT (epithelial to mesenchymal transition) activator *ZEB1* (Supplementary Figure 5C-D) (Burk et al., 2008). *ZEB1* is an LD-TF in soft tissue sarcoma cells (LD-score = 1.5, FDR = 0.007) that are characterized by their mesenchymal origin. We further found that melanoma LD-TFs were differentially expressed in MITF_low_ compared to MITF_high_ samples (Figure 5D). This indicates, that MITF_low_ cells might be able alter their CRC – potentially in order to adapt to the new environment that they invade. We further found that decreased expression of *MITF* was associated with the gain of new gene dependencies. Specifically, dependency on key negative regulators Wnt-signaling (Figure 5E) such as *AXIN1* (FDR = 0.00156), *GSK3B* (FDR = 0.00701) and *APC* (FDR = 0.133) was increased in melanoma cell lines with low *MITF* expression levels (Zhan et al., 2017). Since *MITF* expression is known to be controlled in part by canonical Wnt signaling (Larue and Delmas, 2006), this indicates that destruction complex activity might be required to maintain low levels of *MITF* expression and altered CRC activity.

**Figure 5.**
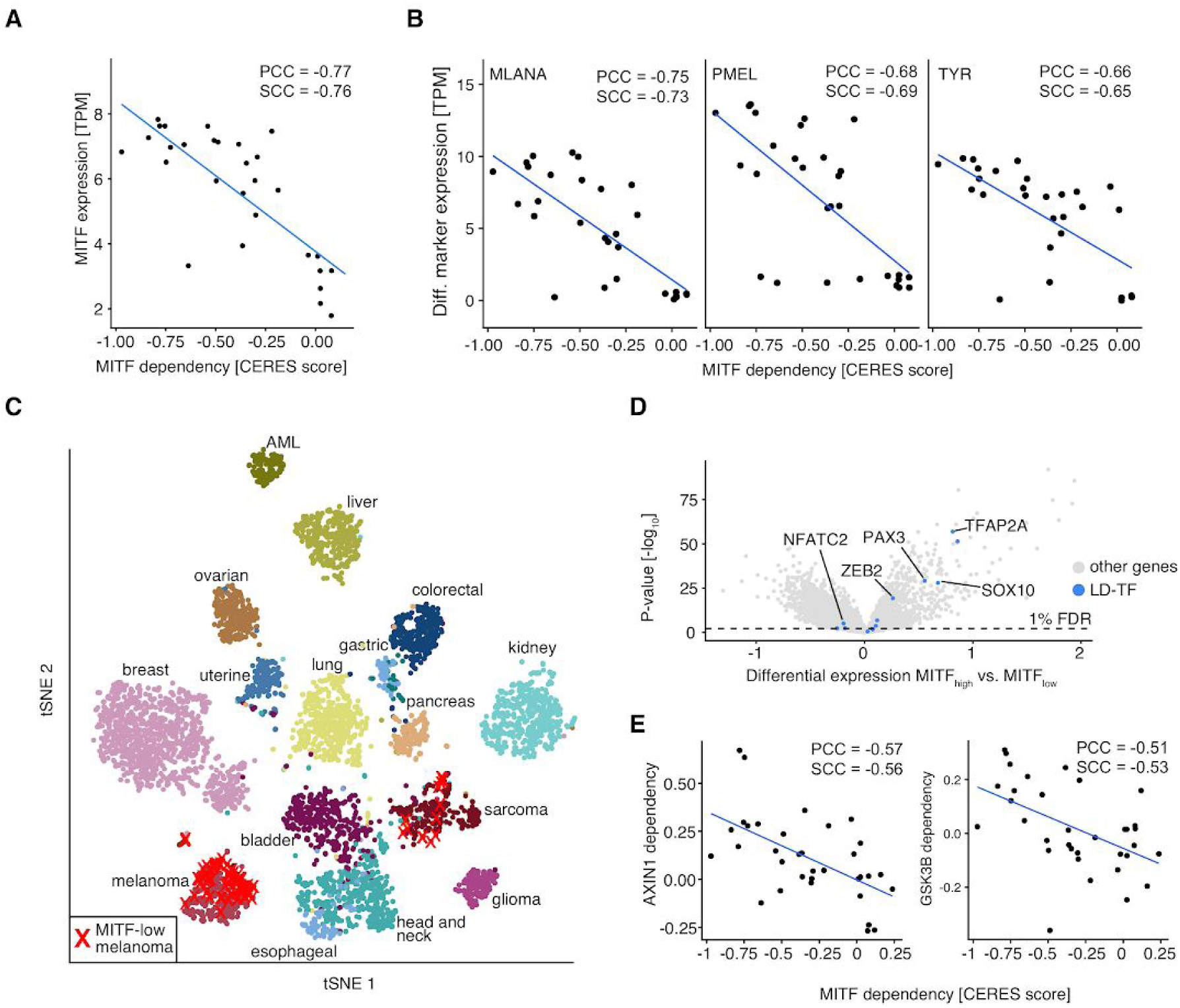
Metastatic cancer cells can alter their core-regulatory circuit to adapt to their new niche. (A) A number of melanoma cell lines are resistant to MITF knockout. This phenomenon is highly correlated with MITF mRNA expression (PCC = Pearson correlation coefficient, SCC = Spearman correlation coefficient). (B) Similarly, loss of MITF expression is correlated with loss of expression of gold-standard melanoma differentiation markers. (C) A t-SNE analysis clusters TCGA tumor samples (n=5,471) by expression of LD-TF genes. Several MITF_low_ melanoma samples (highlighted as red x) cluster among sarcoma tumor samples. (D) A volcano plot highlights differential gene expression in MITF_high_ compared to MITF_low_ patient tumor samples (TCGA; n=367). The strongest melanoma CRC TFs are highlighted as blue dots. (E) Loss of MITF dependency co-occurs with an increased dependency on core-genes of the canonical Wnt-signaling destruction complex such as AXIN1 or GSK3B.

## DISCUSSION

Over the last years large scale efforts have systematically mapped out the essential genome of hundreds of cancer cell lines (Behan et al., 2018; Hart et al., 2015; Meyers et al., 2017; Tsherniak et al., 2017). These studies showed that the majority of essential genes are specific to only a subset of these lines. While some of these context-dependent essential genes could be linked to the presence of certain biomarkers, such as for example somatic mutations in onco- or tumor suppressor genes, the mechanisms that impose selective dependencies remain unknown in most cases. Interestingly, similar context-specificity has been observed for the presence of somatic alterations in cancer driver genes (Haigis et al., 2019).

In this study we integrated genetic dependencies in cancer cells with genomic and transcriptomic data to pinpoint lineage-specific core-regulatory circuits (CRCs) as key drivers of gene essentiality in cancer cells. Building on previous observations in pediatric cancers, we show that selective dependency on CRC-associated transcription factors (TFs) occurs in a diverse range of tumors where they are aberrantly regulated - either as the result of somatic alterations in upstream pathways or variants in the CRC TFs themselves. This effect is not specific to cell lines but is also observed in patient samples. The aberrant regulation of CRC TFs leads to an altered gene expression state that the cells need to maintain in order to maintain a malignant phenotype. As a consequence, tumor cells become dependent on processes that regulate CRC activity as well as key CRC target genes (Figure 6A) (Bradner et al., 2017). We demonstrate how these effects can be exploited to systematically predict CRCs and associated biological processes. We provide lists of predicted CRCs and related pathways for 20 different tumor types as a resource for future study. We propose to use these results as starting points for more in depth characterization of the transcriptional mechanisms that lead to dysregulated CRCs in various cancer types. We further envision, that our results will guide the interpretation of new gene essentiality screens in previously uncharacterized cell lines.

**Figure 6.**
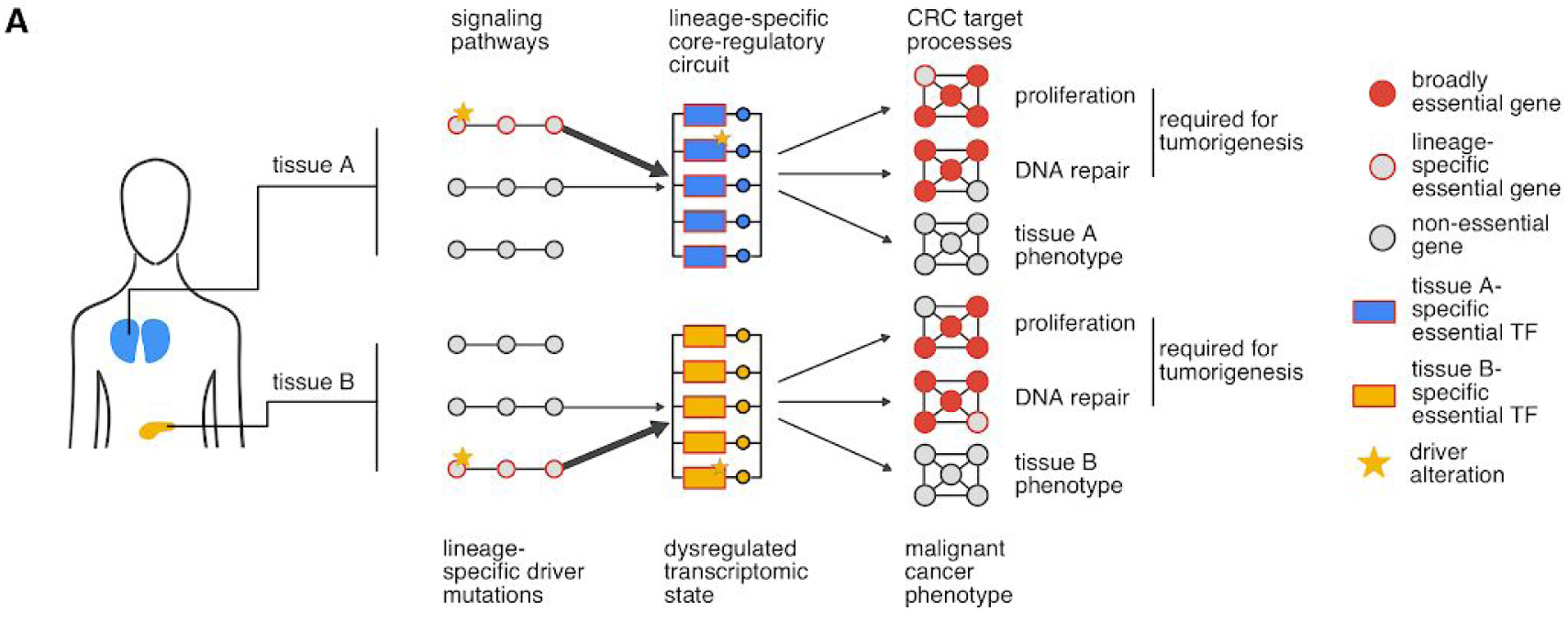
Lineage-specific core-regulatory circuits determine gene essentiality in cancer cells. (A) The transcriptional states of different cell lineages in the human body are controlled by lineage-specific core-regulatory circuits (CRCs). These CRCs consist of a number of transcription factors (TFs) that regulate each other’s activity (center of the diagram). In cancer, CRC activity is dysregulated as the consequence of alterations in upstream signaling pathways or the CRC TFs themselves. This leads to lineage-specific dependency on genes involved in these processes (red border). Depending on CRC composition, the extent to which each pathway contributes to its activity varies. This could explain why most driver alterations themselves are, to some degree, lineage-specific. CRCs control the activity of various biological mechanisms including both general processes (e.g. the cell cycle) and lineage-specific processes that are responsible for the establishment of cell type-specific phenotypes (e.g. melanogenesis in melanocytes). Many genes involved in general processes, that are often required for tumorigenesis, are found to be broadly essential. However, exceptions exist where genes involved in such processes are essential only in certain lineages. In contrast, dependency on genes involved in establishing lineage-specific cell phenotypes is rarely observed in cell lines.

We noticed considerable variation regarding TF dependency in cell lines derived from the same cancer lineages. As an example, we analyze the loss of *MITF* dependency that co-occurs with the loss of *MITF* expression in melanoma cells – a phenomenon termed ‘phenotype switching’ (Li et al., 2015). Our findings indicate that these cells might alter their CRC through changes in signaling pathway activity – perhaps in order to adapt to a change in environment - creating a new set of vulnerabilities. Similar observations have recently been made in Neuroblastoma. There it was shown that two phenotypically distinct differentiation states exist in these tumors that are controlled by different regulatory circuits (van Groningen et al., 2017). This could point to a more general phenomenon where the differentiation state of a tumor cell might affect the set of expressed TFs in a tumor cells which consequently could influence CRC composition. This might also explain the varying number of lineage dependency TFs that we identify for each cancer type. For lung cancer, for example, we found only few TF that showed significant lineage dependency although data from a large number of cell lines was available for this tumor type. In our study we simply stratified cancer cell lines by their tissue of origin. However, it is well known that lung cancers are phenotypically diverse and that these phenotypes are associated with differences in gene expression and response to therapy. It has further been shown that lung adenocarcinomas can adapt their regulatory architecture as they progress by suppressing the core TF *NKX2-1* (Winslow et al., 2011). Since lung cancer cell lines do in fact represent tumors of various phenotypes, we assume that a potential reason for the low number of identified LD-TFs could lie in the way we stratified the cell lines. We thus assume that tissues that are composed of many heterogeneous cell types (controlled by different CRCs) that can each give rise to tumor cells then these might be better examined separately.

In conclusion, our analysis provides the first systematic discovery of CRCs that control the transcriptomic states in different cancer lineages. Specifically, we provide insight into their composition, mechanisms by which they are regulated and their plasticity. Our results confirm that transcriptional addiction is a key driver of selective gene dependency in cancer cells and we show that this effect can be exploited to infer regulatory circuits independent of transcriptomic data. As a platform for future studies we provide a list of predicted CRCs and associated biological processes.

In the future we envision that large-scale single-cell sequencing efforts to map different cell types in the human body (Rozenblatt-Rosen et al., 2017) will provide markers for improved stratification of cell lineages that can give rise to cancer. This might then allow to increase the resolution at which CRCs can be discovered in heterogeneous tissues. Further, CRC TFs are generally characterized by their association with super enhancer elements (Whyte et al., 2013). Super enhancers are usually identified through ChIP-sequencing of H3K27Ac associated genomic regions (Durbin et al., 2018). However, these data exist only for a limited number of cell lines. Additionally, CRC studies might benefit from functional screens with image-based phenotyping where morphological changes upon CRC TF perturbation could be observed. This might provide new insights into the mechanisms by which CRC composition influences cell differentiation.

## METHODS

### Context-specific gene dependencies

To determine context-specific gene dependencies, Avana CRISPR fitness screens were downloaded from the Cancer Dependency Map (DepMap) portal (Supplementary Table 3). A two component Gaussian mixture model was then fit to the CERES scores (Meyers et al., 2017) measured across all 558 cancer cell lines for each gene in the CRISPR library using the ‘mixtools’ R package (Benaglia et al., 2009). Each data point was assigned to one of the mixture components by their posterior probability and the component representing the majority of data was selected as the baseline phenotype distribution of a gene. This is based on the assumption that a particular gene knockout has a similar effect on the majority of cell lines while only a smaller subset of cell lines are especially vulnerable (context-dependent) or resistant to that knockout (Costanzo et al., 2010, 2016). For a few genes (78 out of 17,634) a two component mixture could not be fit successfully due to high uniformity of knockout phenotypes across cell lines. These were excluded from downstream analyses. Also, the gene SOX9 was excluded due to known off-target effects of sgRNAs in the Avana library (Fortin et al., 2019). Next, a p-value was computed for each knockout phenotype in each gene representing the probability of having observed that phenotype under the baseline phenotype distribution of that gene. Specifically, we tested the null hypothesis *H*_0_: μ_*g*_ = *y*_*cg*_ where μ_*g*_ is the mean of the baseline phenotype distribution for gene *g* (as determined above) and *y*_*cg*_ is the CERES score measured in cell line *c* upon knockout of gene *g*. Resulting p-values were adjusted for multiple testing using the Benjamini-Hochberg method and context-dependent vulnerabilities were determined at 20% false discovery rate (FDR). Gene set enrichment analysis was performed on all genes ranked by the number of context-dependent cell lines using the ‘fgsea’ R/Bioconductor (Gentleman et al., 2004; Huber et al., 2015; Sergushichev, 2016) package on Gene Ontology (Gene Ontology Consortium, 2015) Molecular Function gene sets of size 15-500 using 100,000 permutations. GO terms and annotations were downloaded through the ‘GO.db’ and ‘Organism.dplyr’ R/Bioconductor packages. Transcription factor (TF) status was assigned for each gene according to Lambert et al (Lambert et al., 2018). For clustering of cell lines by context-specific TF genes, TF for which context-dependency was determined in more than 25 different cancer cell lines were selected and CERES scores for each TF were centered by subtracting the mean of their baseline phenotype distribution μ_*g*_.

### Calculation of lineage dependency scores (LD-scores)

For LD-score calculation, all cancer lineages for which CRISPR screening data in more than 10 different cell lines was available were retained (in total 20 lineages). LD-scores where then inferred for each gene via a linear model as

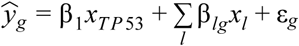

Where 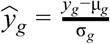 are the CERES scores measured for gene *g* after centering by subtraction of the mean and scaling through division by the standard deviation of the baseline phenotype distribution for gene *g*. Further, *x*_*TP* 53_ represents the *TP53* mutation status and *x*_*l*_ are the cancer lineages for each cell line. β_*lg*_ are then lineage dependency scores (LD-scores) for each gene in each cancer lineage. P-values for all LD-scores were adjusted for multiple testing using the Benjamini-Hochberg method. *TP53* mutation status was determined for each cell line according to Cancer Cell Line Encyclopedia and DepMap cell line mutation data (Barretina et al., 2012). *TP53* mutations were considered if they were flagged as either ‘isDeleterious’ or ‘isTCGAhotspot’ in the DepMap data set. Silent mutations were excluded. It has been shown that a cell line’s *TP53* mutation status has a major impact on its gene essentiality profile independent of the cell line’s tissue of origin (Rauscher et al., 2018; Tsherniak et al., 2017). Therefore, we decided to account for *TP53* mutations when calculating LD-scores.

### Assembly of putative core-regulatory circuits

First, significant lineage dependency TFs were selected (LD-score > 1, FDR < 0.05). To avoid false positives due to potential CRISPR off-target effects, candidate LD-TF not expressed in their associated cancer lineage (median TPM across cell lines of corresponding lineage < 0.1) or reported as loss-of-function tolerant in ExAC (LOF Z-score < 1) were excluded from downstream analyses. Both thresholds were informed by manual examination of known non-expressed/non-essential genes. To determine whether TF target gene-relationships occur at increased frequency between TF that act in the same CRCs, TF-target gene annotations were downloaded from the Harmonizome portal (Rouillard et al., 2016). Specifically, we downloaded the CHEA, ENCODE, ESCAPE and manually curated TRANSFAC annotation lists (ENCODE Project Consortium, 2004; Lachmann et al., 2010; Matys et al., 2003; Xu et al., 2013). For each cancer lineage we then counted the number of unique relationships between pairs of significant LD-TFs reported in the above data sets. To understand whether the number of relationships observed between the LD-TFs associated with a cancer lineage was higher than expected by chance, we randomly sampled TFs included in the Harmonizome data and calculated the number of relationships reported between these to infer a null distribution. A Kolmogorov-Smirnov test was then used to compute the probability of having observed the number of regulatory relationships between LD-TFs genes by chance. To test whether LD-TFs are frequently linked to super enhancer (SE) elements in their associated lineages, SE data were downloaded from SEdb (Jiang et al., 2018) for all samples that could be matched to one of the lineages represented by the CRISPR data. Supplementary Table 3 contains SEdb identifiers for samples used for analysis. Each SE element was linked to its closest active gene on the genome as defined by SEdb. Processed HCT116 ChIP-seq peaks were downloaded for H3K27ac, POLR2A and TCF7L2 targets from ENCODE (ENCODE Project Consortium, 2004).

### Molecular characterization of lineage dependency transcription factors

First, genes were divided into significant lineage dependency (LD) TF (LD-score > 1 and FDR < 5%), other TF and non-TF genes. Functional gene constraint scores were downloaded for non-cancer samples from ExAC (Supplementary Table 3) (Lek et al., 2016). Between the above gene groups, we compared (i) ‘LOF Z-scores’ which quantify whether a damaging mutation in a gene is observed more or less frequently than expected by chance and (ii) ‘pLI scores’ scores, which represent the probability of a gene being loss-of-function intolerant. To determine whether LD-TFs are expressed at higher levels in associated normal tissues compared to other tissues, normalized GTEx RNA-seq gene expression values (TPM) were downloaded (GTEx Consortium et al., 2017). Where possible, GTEx tissues were matched to cancer cell line lineages (Supplementary Table 4). In total, 17 lineages could be matched. Transformed fibroblast and lymphocyte samples were excluded. To enable comparison of expression levels between different genes, log(TPM+1) values for each gene were scaled to range [0,1] where 0 represents no/minimal gene expression and 1 indicates the maximal expression observed across all processed samples. To test whether LD-TF expression is increased in cancer samples of the associated tumor types, normalized RNA-seq data were downloaded from TCGA (Cancer Genome Atlas Research Network et al., 2013) using the ‘RTCGA’ R/Bioconductor package (Kosinski et al., 2016) and TCGA tumor type annotations were matched to cancer cell line tumor types (Supplementary Table 4). Similar to the GTEx data, expression values for each gene were scaled to range [0,1] before comparison. To test whether LD-TF are differentially regulated in cancer compared to normal tissue samples, TCGA RNA-seq data were selected for all tumor types where corresponding normal sample data was available (12 in total, Supplementary Table 4). Differential gene expression between healthy and tumor samples was determined for each LD-TF associated with one of the 12 cancer types (in total 158) using a two-sample Welch’s t-test. The Benjamini-Hochberg method was used to control the false discovery rate. Each LD-TF was then categorized as downregulated (μ(*T*) - μ(*H*) > 0.1 and *FDR* < 20%), upregulated (μ(*H*) - μ(*T*) < −0.1 and *FDR* < 20%) or not differentially regulated where μ(*T*) is the average expression of the gene in log(FPKM+1) in tumor samples and μ(*H*) is the average expression in healthy samples. A similar analysis was performed for 1,000 equally large sets of randomly selected genes to infer a distribution that indicates how often up- or down-regulation would be expected by chance.

To determine whether LD-TFs are affected by somatic alterations in patient cancer samples, mutation, copy number alteration, methylation and gene fusion data were downloaded from the COSMIC database (Supplementary Table 3) (Forbes et al., 2017). Silent mutations were excluded. Additionally, gene fusions, copy number gains and hypomethylation events were selected for each sample as reported by COSMIC. If a specific type of variant was observed for a gene in more than 5% of all samples from a certain tumor type, that gene was considered as frequently altered. A Fisher’s exact test was performed to determine whether alterations in LD-TF occur more frequently compared to other genes.

### Codependency analysis of lineage dependency genes

Lineage dependency scores were calculated for all non-transcription factor genes as described above. Next, significant LD-genes were selected for each cancer lineage (LD-score > 1, FDR < 5%). Genes were excluded if their median expression in cell lines of their associated tissues was below 0.01 (TPM). Gene codependency was then calculated for all possible gene pairs as the Pearson correlation coefficient of their CERES scores across all 558 cell lines in the DepMap data set. LD-genes were clustered into groups according to their codependency using a model based clustering approach implemented in the R package ‘mclust’. Here, Gaussian mixture models with up to 10 components (potential clusters) are fitted to the data and the ideal number of clusters is selected according to the Bayesian Information Criterion (BIC) for each model. The maximal number of clusters selected for any lineage was 9 indicating that allowing for up to 10 clusters was appropriate. Next, gene-set enrichment analysis was performed to identify significantly overrepresented Gene Ontology (Gene Ontology Consortium, 2015) Biological Process terms for each cluster. To this end, the methods ‘goana’ and ‘topGO’ implemented in the ‘limma’ R/Bioconductor package were used (Alexa and Rahnenfuhrer, 2010; Ritchie et al., 2015).

### MITF-dependency in melanoma

Normalized gene expression data for melanoma patient samples were downloaded from TCGA using the Rtcga R/Bioconductor package (367 samples) (Huber et al., 2015; Kosinski et al., 2016). Differentially expressed genes between MITF_high_ and MITF_low_ samples were determined as the degree of coexpression between *MITF* and all other genes using the linear model *y*_*g*_ = β_0_ + β_*MITF*_ *x*_*MITF*_ + ε_*g*_ for all genes *g* where *y*_*g*_ is the gene expression of gene *g* and *x*_*MITF*_ is the gene expression of *MITF* across all melanoma patient samples. The coefficient β_*MITF*_ quantifies the degree of coexpression between *MITF* and gene *g*. The Benjamini-Hochberg method was used to control the false discovery rate (FDR) at 1%. *MITF* codependency in melanoma cancer cell lines (34 cell lines) was determined in a similar manner using CRISPR knockout phenotypes (CERES scores) instead of gene expression values. T-distributed stochastic neighbor embedding (t-SNE; as implemented in the ‘Rtsne’ R-package) was performed to cluster tumor samples based on gene expression of significant (LD-score > 1, FDR < 5%) lineage dependency transcription factors.

## Data and Software Availability

Computer code to reproduce all figures in the manuscript is available online at: https://github.com/boutroslab/Supplemental-Material/tree/master/Rauscher_2019.

## ACKNOWLEDGEMENTS

We would like to thank the Cancer Dependency Map (DepMap) Consortium that provided CRISPR fitness screening data for analysis. We thank M. Funk for providing illustrations. We are grateful to members of the Boutros lab for comments on the manuscript and valuable discussion. B.R. was supported by the BMBF-funded Heidelberg Center for Human Bioinformatics (HD-HuB) within the German Network for Bioinformatics Infrastructure (de.NBI). Work in the Boutros lab is supported in part by a grant from the European Research Council (ERC).

