## Supplementary Information for "Lineage specific core-regulatory circuits determine gene essentiality in cancer cells"

Rauscher et al.

**Contents**

1. **Supplementary Figures**
2. **Supplementary Tables**

### Supplementary Figures

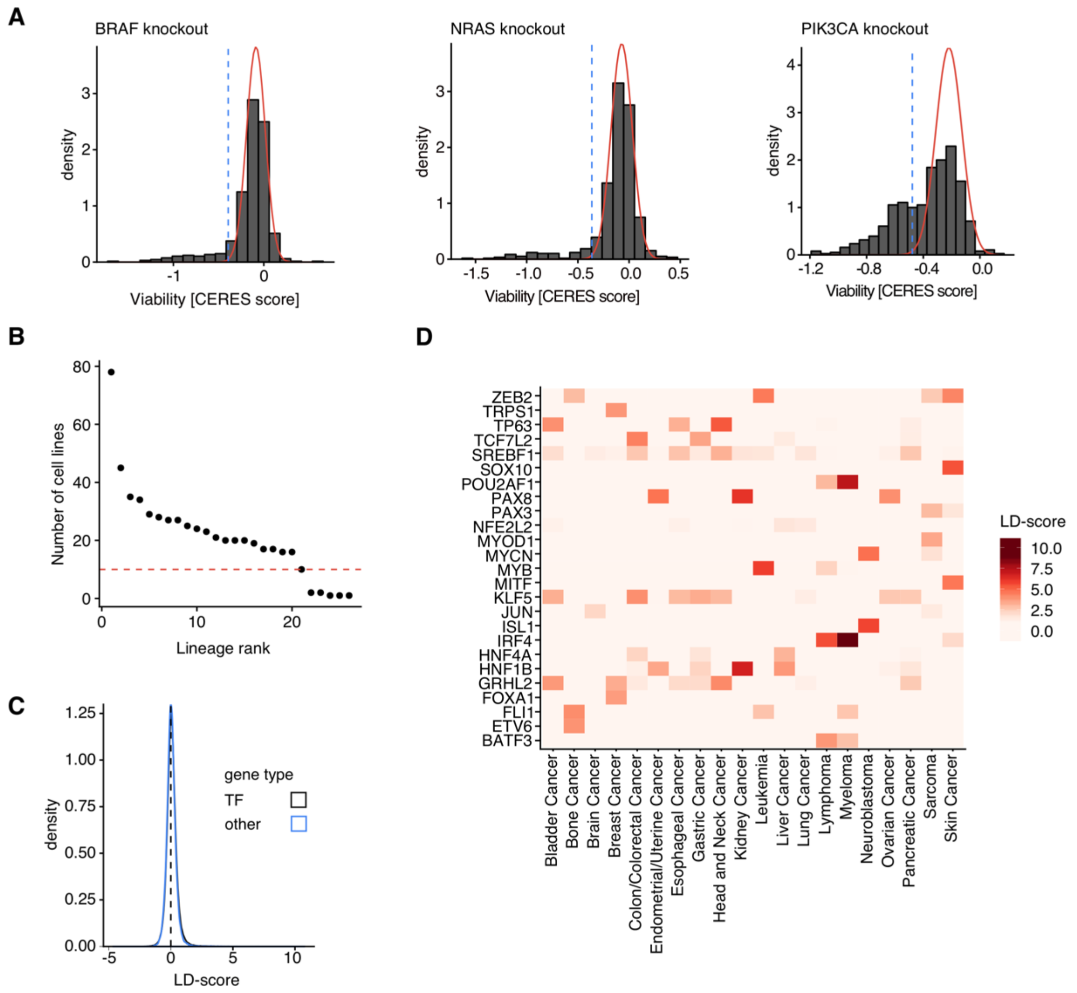


**Supplementary Figure 1.** (A) Histograms of CERES scores measured upon knockout of BRAF, NRAS and PIK3CA illustrate context-specific gene dependency in cases of oncogene addiction. The red curve indicates the estimated baseline phenotype. The vertical blue line marks a 20% FDR cutoff separating expected from context-specific phenotypes. (B) Number of cell lines available for each cancer lineage in the DepMap 19q1 dataset. Each dot represents one linage and the lineages are ranked by the number of available cell lines. The red horizontal line highlights the n=10 cutoff. (C) Density plot of inferred LD-scores for all genes. The black curve represents transcription factors (TF) while the blue curve represents all other genes. (D) A heatmap of LD-scores for the 2 highest scoring TFs in each cancer lineage tested.

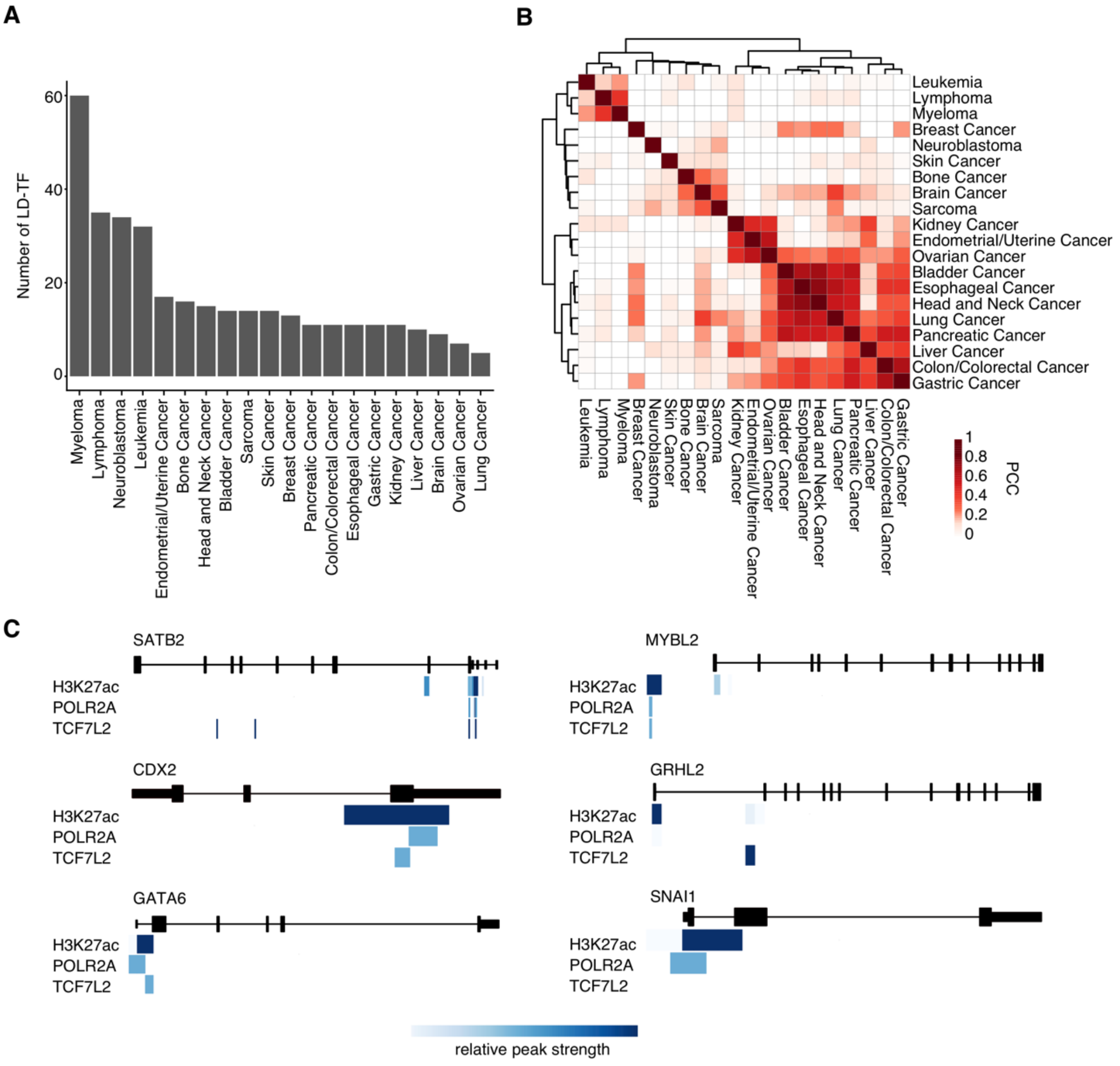


**Supplementary Figure 2.** (A) Number of significant LD-TFs (LD-score > 1, FDR < 5%) identified for each cancer lineage. (B) Clustering of cancer lineages by the Pearson correlation coefficient (PCC) of their LD-score profiles for all transcription factors. (C) Public ChIP-seq data on H3K27ac, POLR2A and TCF7L2 target proteins shows POLR2A and TCF7L2 binding in enhancer regions that are found close to the transcription start sites of colorectal cancer CRC genes. Blue color intensity represents the relative peak strength. No peaks for found at the *HNF1A* locus.

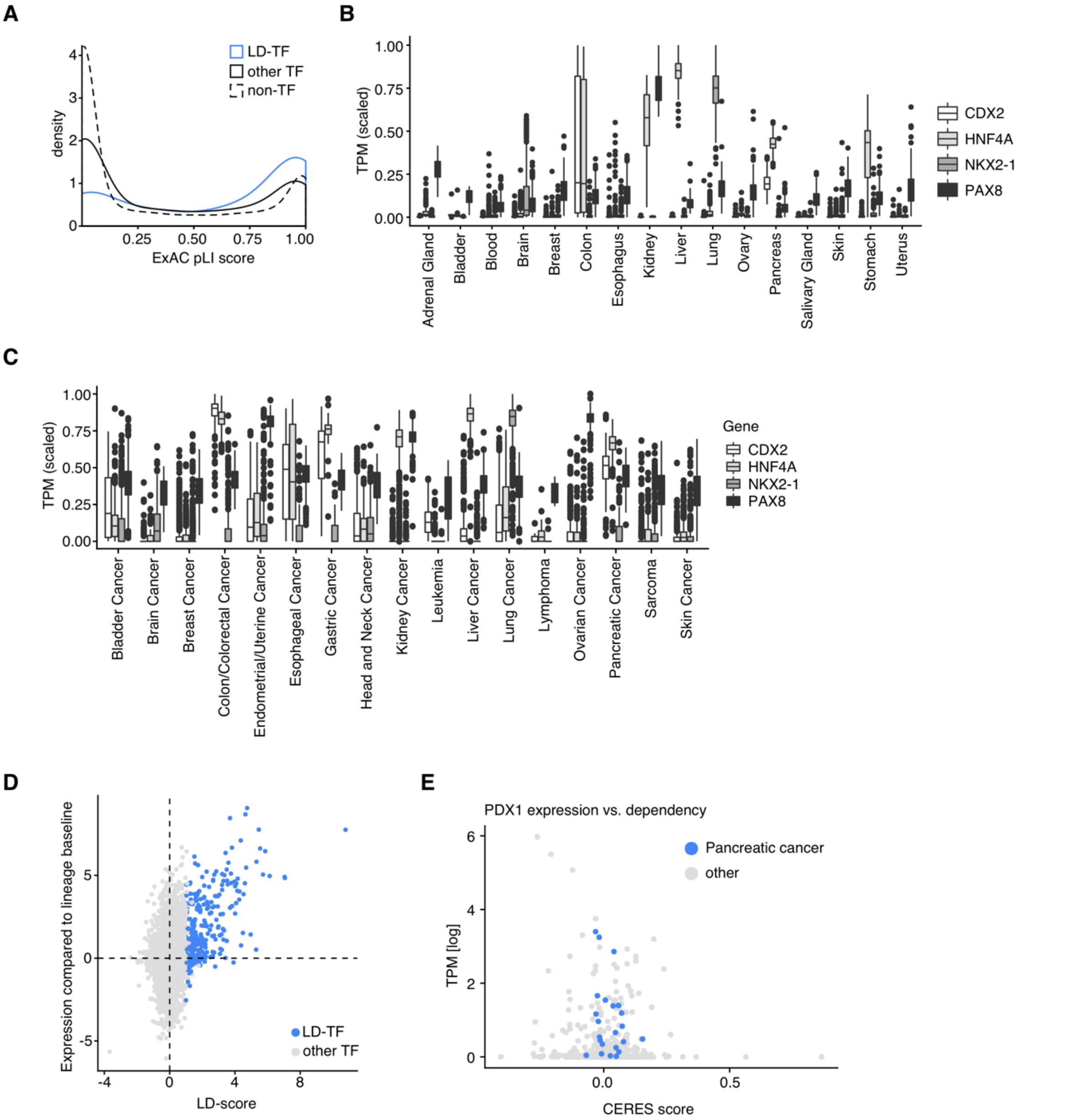


**Supplementary Figure 3.** (A) pLI-score distributions for significant LD-TFs (LD-score >1, FDR < 5%), other TFs and non-TF genes in the ExAC database. pLI represents the probability of a gene being loss-of-function intolerant (intolerant of both heterozygous and homozygous loss-of-function variants). (B) Scaled expression values for selected transcription factor genes in different tissues represented in the GTEx dataset. The center of each box is the sample median, the whiskers extend from the upper (lower) hinge to the largest (smallest) data point no further than 1.5 times the interquantile range from the upper (lower) hinge. (C) Scaled gene expression values for selected TF genes in different cancer types represented by the TCGA dataset. (D) Comparison of LD-scores and corresponding gene expression values of transcription factor genes in cancer cell lines. Each dot represents a unique combination of TF and cancer lineage. Blue dots represent significant LD-TFs (LD-score > 1, FDR < 5%). (E) Comparison of gene essentiality (CERES score) and gene expression of the pancreatic and duodenal homeobox 1 (*PDX1*) gene. Each dot represents a cancer cell line, pancreatic cell lines are highlighted in blue.

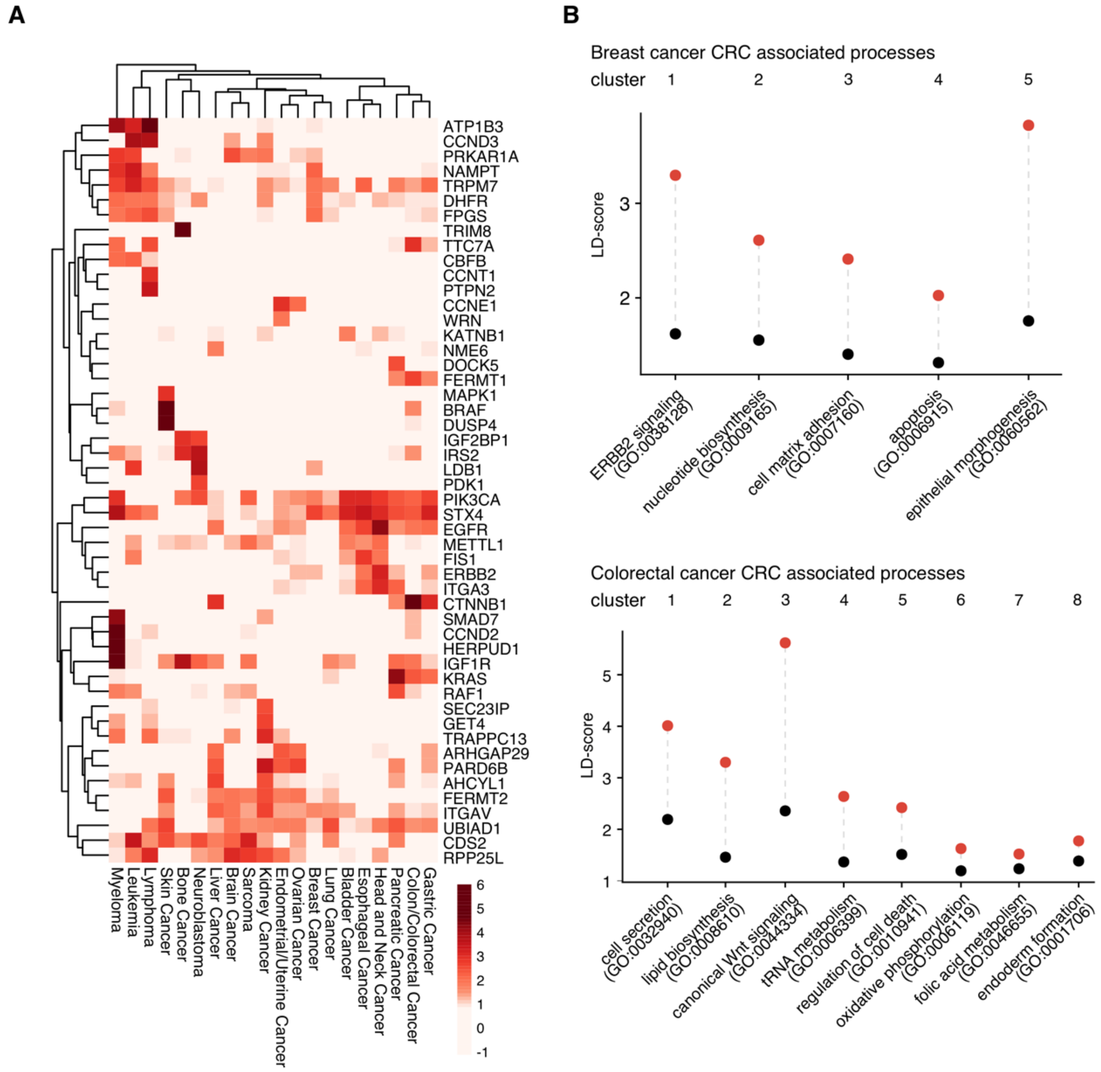


**Supplementary Figure 4.** (A) A heatmap of LD-scores of the 3 strongest (by LD-score) lineage-specific non-transcription factor dependencies for each cancer type tested. (B) Biological processes associated with the core-regulatory circuits of breast cancer and colorectal cancer. Dots indicate average (black dot) and maximum (red dot) LD-scores for genes associated with each codependency cluster of lineage dependency genes. A representative enriched GO Molecular Function term is shown for each cluster.

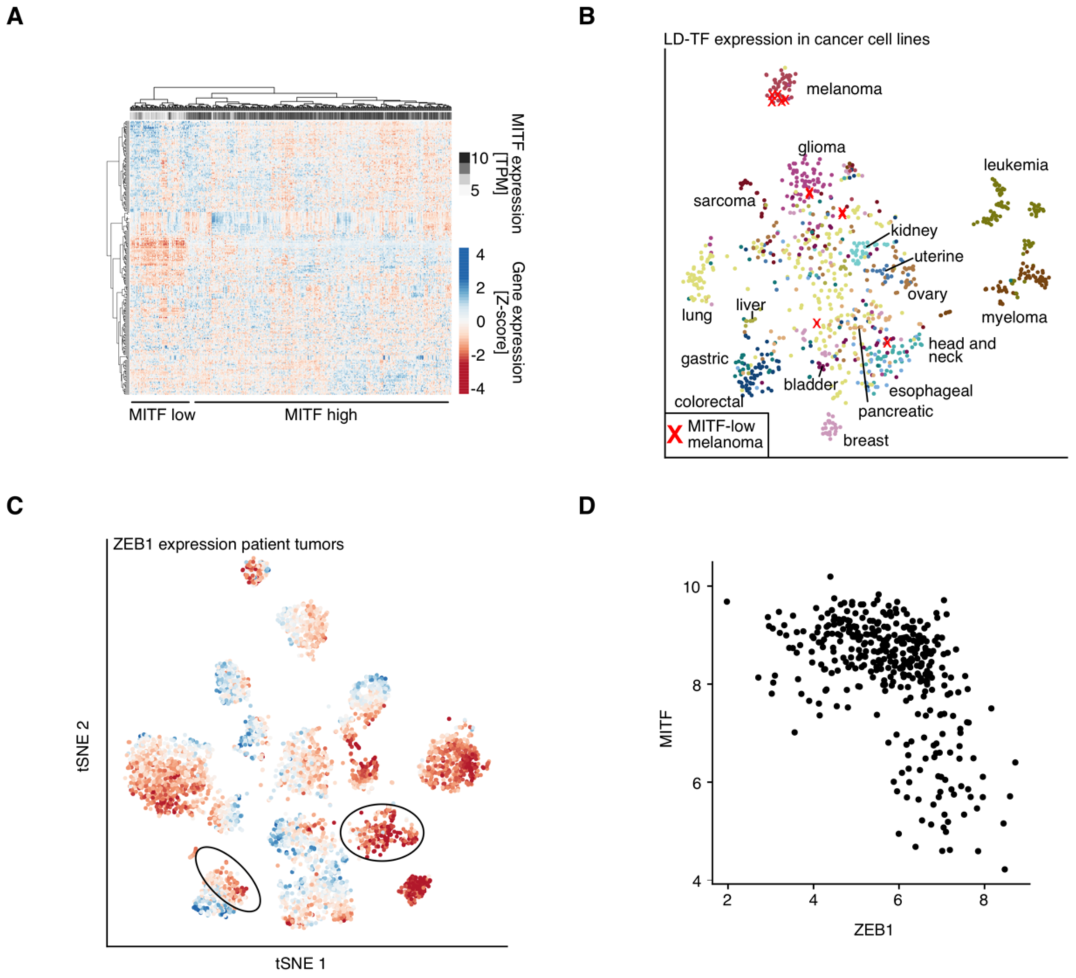


**Supplementary Figure 5.** (A) A heatmap of scaled gene expression values of 250 genes with the highest expression standard deviation across melanoma samples (y-axis) in TCGA melanoma patient tumor samples. The annotation bar at the top indicates the *MITF* expression level for each sample. (B) t-SNE analysis clusters cancer cell lines by expression of LD-TF genes. $MITF_{low}$ melanoma cell lines are highlighted as red x. (C) t-SNE analysis of TCGA patient tumor samples visualizes the relative gene expression of the ZEB1 gene across tumor samples. Blue color represents low levels of gene expression and red color indicates high expression of *ZEB1*. Black circles highlight regions where $MITF_{low}$ samples are situated in the plot. (D) Expression levels of MITF compared to ZEB1 in melanoma patient samples (n=367). Each dot represents one tumor sample. Gene expression values are log(FPKM + 1).

### Supplementary Tables

| Dataset | File name | Access date | Link |
|---|---|---|---|
| DepMap CRISPR screens | DepMap-2019q1-celllines.csv | February 22nd, 2019 | https://depmap.org/portal/download/ |
| DepMap cell line annotation | DepMap-2019-q1-celllines.csv | February 22nd, 2019 | https://depmap.org/portal/download/ |
| Human Transcription Factors | DatabaseExtract_v_1.01 | April 9th, 2019 | http://humantfs.ccbr.utoronto.ca/download.php |
| DepMap mutation calls | Depmap_19Q1_mutation_calls.csv | February, 22nd, 2019 | https://depmap.org/portal/download/ |
| DepMap/CCLE gene expression | CCLE_18q4_TPM.csv | December 21st, 2019 | https://depmap.org/portal/download/ |
| DEMETER2 RNAi screens | D2_combined_gene_dep_scores.csv | February, 22nd, 2019 | https://depmap.org/portal/download/ |
| Harmonizome ENCODE TF targets | Gene_attribute_matrix.txt.gz | February, 26th, 2019 | http://amp.pharm.mssm.edu/Harmonizome/dataset/ENCODE+Transcription+Factor+Targets |
| Harmonizome CHEA TF targets | Gene_attribute_matrix.txt.gz | February, 26th, 2019 | http://amp.pharm.mssm.edu/Harmonizome/dataset/CHEA+Transcription+Factor+Targets |
| Harmonizome TRANSFAC TF targets | Gene_attribute_matrix.txt.gz | February, 26th, 2019 | http://amp.pharm.mssm.edu/Harmonizome/dataset/TRANSFAC+Predicted+Transcription+Factor+Targets |
| Harmonizome ESCAPE TF targets | Gene_attribute_matrix.txt.gz | February, 26th, 2019 | http://amp.pharm.mssm.edu/Harmonizome/dataset/ESCAPE+Omics+Signatures+of+Genes+and+Proteins+for+Stem+Cells |
| SEdb super enhancer data | See below for details | March 21st, 2019 | http://www.licpathway.net/sedb/download.php |
| HCT116 ChIP-sequencing | ENCFF333WEH.bed, ENCFF804JZT.bed, ENCFF083XMT.bed | March 11th, 2019 | https://www.encodeproject.org/search/?type=Experiment&status=released |
| ExAC non-cancer data | fordist_cleaned_exac_nonTCGA_z_pli_rec_null_data.txt | February 23rd, 2019 | ftp://ftp.broadinstitute.org/pub/ExAC_release/current/functional_gene_constraint |
| GTEx expression data | GTEx_Analysis_2016-01-15_v7_RNASeQCv1.1.8_g | February 23rd, 2019 | https://gtexportal.org/home/datasets |

| | | | |
|---|---|---|---|
| | ene_tpm.gct.gz | | |
| GTEx sample information | GTEx_v7_Annotations_SampleAttributesDS.txt | February 23rd, 2019 | https://gtexportal.org/home/datasets |
| TCGA gene expression data | RTCGA.rnaseq R package | Version 20151101.8.0 | https://www.bioconductor.org/packages/release/data/experiment/html/RTCGA.rnaseq.html |
| COSMIC mutations | CosmicGenomeScreensMutantExport.tsv.gz | February 23rd, 2019 | https://cancer.sanger.ac.uk/cosmic/download |
| COSMIC copy numbers | CosmicCompleteCNA.tsv.gz | February 23rd, 2019 | https://cancer.sanger.ac.uk/cosmic/download |
| COSMIC gene fusions | CosmicFusionExport.tsv.gz | February 23rd, 2019 | https://cancer.sanger.ac.uk/cosmic/download |
| COSMIC methylation | CosmicCompleteDifferentialMethylation.tsv.gz | February 23rd, 2019 | https://cancer.sanger.ac.uk/cosmic/download |

**Supplementary Table 3: Used datasets**

List of SEdb files used:
Sample_00_005_SE.bed, Sample_00_006_SE.bed, Sample_00_007_SE.bed, Sample_00_008_SE.bed, Sample_00_009_SE.bed, Sample_00_011_SE.bed, Sample_00_015_SE.bed, Sample_00_020_SE.bed, Sample_00_029_SE.bed, Sample_00_030_SE.bed, Sample_00_035_SE.bed, Sample_00_041_SE.bed, Sample_00_042_SE.bed, Sample_00_043_SE.bed, Sample_01_006_SE.bed, Sample_01_012_SE.bed, Sample_01_020_SE.bed, Sample_01_024_SE.bed, Sample_01_025_SE.bed, Sample_01_030_SE.bed, Sample_01_034_SE.bed, Sample_01_037_SE.bed, Sample_01_038_SE.bed, Sample_01_043_SE.bed, Sample_01_046_SE.bed, Sample_01_048_SE.bed, Sample_01_052_SE.bed, Sample_01_056_SE.bed, Sample_01_057_SE.bed, Sample_01_059_SE.bed, Sample_01_071_SE.bed, Sample_01_074_SE.bed, Sample_01_075_SE.bed, Sample_01_077_SE.bed, Sample_01_082_SE.bed, Sample_01_083_SE.bed, Sample_01_089_SE.bed, Sample_02_002_SE.bed, Sample_02_003_SE.bed, Sample_02_004_SE.bed, Sample_02_008_SE.bed, Sample_02_009_SE.bed, Sample_02_012_SE.bed, Sample_02_013_SE.bed, Sample_02_014_SE.bed, Sample_02_015_SE.bed, Sample_02_016_SE.bed, Sample_02_021_SE.bed, Sample_02_022_SE.bed, Sample_02_023_SE.bed, Sample_02_025_SE.bed, Sample_02_026_SE.bed, Sample_02_029_SE.bed, Sample_02_030_SE.bed, Sample_02_031_SE.bed, Sample_02_032_SE.bed, Sample_02_033_SE.bed, Sample_02_034_SE.bed, Sample_02_035_SE.bed, Sample_02_051_SE.bed, Sample_02_052_SE.bed, Sample_02_055_SE.bed, Sample_02_056_SE.bed, Sample_02_058_SE.bed, Sample_02_060_SE.bed, Sample_02_069_SE.bed, Sample_02_070_SE.bed, Sample_02_071_SE.bed, Sample_02_072_SE.bed, Sample_02_077_SE.bed, Sample_02_078_SE.bed, Sample_02_079_SE.bed, Sample_02_081_SE.bed, Sample_02_086_SE.bed, Sample_02_087_SE.bed,

Sample_02_095_SE.bed, Sample_02_096_SE.bed, Sample_02_100_SE.bed,
Sample_02_101_SE.bed, Sample_02_102_SE.bed, Sample_02_103_SE.bed,
Sample_02_104_SE.bed, Sample_02_110_SE.bed, Sample_02_111_SE.bed,
Sample_02_114_SE.bed, Sample_02_115_SE.bed, Sample_02_116_SE.bed,
Sample_02_126_SE.bed, Sample_02_127_SE.bed, Sample_02_131_SE.bed,
Sample_02_132_SE.bed, Sample_02_134_SE.bed, Sample_02_135_SE.bed,
Sample_02_136_SE.bed, Sample_02_138_SE.bed, Sample_02_139_SE.bed,
Sample_02_140_SE.bed, Sample_02_141_SE.bed, Sample_02_142_SE.bed,
Sample_02_143_SE.bed, Sample_02_144_SE.bed, Sample_02_147_SE.bed,
Sample_02_148_SE.bed, Sample_02_150_SE.bed, Sample_02_151_SE.bed,
Sample_02_152_SE.bed, Sample_02_161_SE.bed, Sample_02_164_SE.bed,
Sample_02_165_SE.bed, Sample_02_166_SE.bed, Sample_02_171_SE.bed,
Sample_02_174_SE.bed, Sample_02_175_SE.bed, Sample_02_177_SE.bed,
Sample_02_179_SE.bed, Sample_02_180_SE.bed, Sample_02_181_SE.bed,
Sample_02_182_SE.bed, Sample_02_186_SE.bed, Sample_02_187_SE.bed,
Sample_02_188_SE.bed, Sample_02_189_SE.bed, Sample_02_190_SE.bed,
Sample_02_191_SE.bed, Sample_02_192_SE.bed, Sample_02_193_SE.bed,
Sample_02_194_SE.bed, Sample_02_195_SE.bed, Sample_02_196_SE.bed,
Sample_02_197_SE.bed, Sample_02_198_SE.bed, Sample_02_199_SE.bed,
Sample_02_200_SE.bed, Sample_02_201_SE.bed, Sample_02_202_SE.bed,
Sample_02_203_SE.bed, Sample_02_204_SE.bed, Sample_02_205_SE.bed,
Sample_02_206_SE.bed, Sample_02_207_SE.bed, Sample_02_208_SE.bed,
Sample_02_209_SE.bed, Sample_02_217_SE.bed, Sample_02_218_SE.bed,
Sample_02_220_SE.bed, Sample_02_226_SE.bed, Sample_02_227_SE.bed,
Sample_02_228_SE.bed, Sample_02_235_SE.bed, Sample_02_236_SE.bed,
Sample_02_237_SE.bed, Sample_02_240_SE.bed, Sample_02_241_SE.bed,
Sample_02_242_SE.bed, Sample_02_246_SE.bed, Sample_02_248_SE.bed,
Sample_02_249_SE.bed, Sample_02_250_SE.bed, Sample_02_252_SE.bed,
Sample_02_253_SE.bed, Sample_02_254_SE.bed, Sample_02_258_SE.bed,
Sample_02_267_SE.bed, Sample_02_268_SE.bed, Sample_02_269_SE.bed,
Sample_02_270_SE.bed, Sample_02_271_SE.bed, Sample_02_272_SE.bed,
Sample_02_280_SE.bed, Sample_02_281_SE.bed, Sample_02_282_SE.bed,
Sample_02_283_SE.bed, Sample_02_285_SE.bed, Sample_02_286_SE.bed,
Sample_02_295_SE.bed, Sample_02_298_SE.bed, Sample_02_299_SE.bed,
Sample_02_300_SE.bed, Sample_02_301_SE.bed, Sample_02_302_SE.bed,
Sample_02_303_SE.bed, Sample_02_304_SE.bed, Sample_02_305_SE.bed,
Sample_02_307_SE.bed, Sample_02_308_SE.bed, Sample_02_309_SE.bed,
Sample_02_310_SE.bed, Sample_02_311_SE.bed, Sample_02_312_SE.bed,
Sample_02_313_SE.bed, Sample_02_314_SE.bed, Sample_02_315_SE.bed,
Sample_02_316_SE.bed, Sample_02_317_SE.bed, Sample_02_318_SE.bed,
Sample_02_319_SE.bed, Sample_02_320_SE.bed, Sample_02_321_SE.bed,
Sample_02_322_SE.bed, Sample_02_323_SE.bed, Sample_02_324_SE.bed,
Sample_02_325_SE.bed, Sample_02_326_SE.bed, Sample_02_327_SE.bed,
Sample_02_338_SE.bed, Sample_02_339_SE.bed, Sample_02_340_SE.bed,
Sample_02_342_SE.bed, Sample_02_344_SE.bed, Sample_02_345_SE.bed,
Sample_02_346_SE.bed, Sample_02_347_SE.bed, Sample_02_348_SE.bed,
Sample_02_349_SE.bed, Sample_02_352_SE.bed, Sample_02_353_SE.bed,
Sample_02_354_SE.bed, Sample_02_355_SE.bed, Sample_02_356_SE.bed,
Sample_02_357_SE.bed, Sample_02_358_SE.bed, Sample_02_359_SE.bed,
Sample_02_361_SE.bed, Sample_02_379_SE.bed, Sample_02_383_SE.bed,
Sample_02_384_SE.bed, Sample_02_389_SE.bed, Sample_02_390_SE.bed,
Sample_02_391_SE.bed, Sample_02_393_SE.bed, Sample_02_394_SE.bed,
Sample_02_397_SE.bed, Sample_02_398_SE.bed, Sample_02_399_SE.bed,
Sample_02_400_SE.bed, Sample_02_401_SE.bed, Sample_02_402_SE.bed,

Sample_02_403_SE.bed, Sample_02_404_SE.bed, Sample_02_405_SE.bed, Sample_02_406_SE.bed, Sample_02_407_SE.bed, Sample_02_408_SE.bed, Sample_02_409_SE.bed, Sample_02_410_SE.bed, Sample_02_411_SE.bed, Sample_02_412_SE.bed, Sample_02_413_SE.bed, Sample_02_415_SE.bed, Sample_02_421_SE.bed, Sample_02_424_SE.bed, Sample_02_426_SE.bed, Sample_02_427_SE.bed, Sample_02_428_SE.bed, Sample_02_429_SE.bed, Sample_02_430_SE.bed, Sample_02_433_SE.bed, Sample_02_434_SE.bed, Sample_02_435_SE.bed, Sample_02_436_SE.bed, Sample_03_004_SE.bed, Sample_03_005_SE.bed, Sample_03_007_SE.bed

**Tissue mapping between resources**

| DepMap (Primary Disease) | GTEx (SMTSD) | TCGA (Project ID) | COSMIC (Primary site) |
|---|---|---|---|
| Bladder Cancer | Bladder (n=11) | BLCA (n=427) | Urinary_tract (n=385) |
| Bone Cancer | | | Bone (n=414) |
| Brain Cancer | Brain (n=2076) | GBM (n=165) | Central_nervous_system (n=1841) |
| Breast Cancer | Breast (n=306) | BRCA (n=1202) | Breast (n=2334) |
| Colon/Colorectal Cancer | Colon (n=539) | COAD (n=328) | Large_intestine (n=2132) |
| Endometrial/Uterine Cancer | Uterus (n=117) | UCEC (n=199) | Endometrium (n=305) |
| Esophageal Cancer | Esophagus (n=1111) | ESCA (n=171) | Oesophagus (n=1078) |
| Gastric Cancer | Stomach (n=272) | STAD (n=34) | Stomach (n=637) |
| Head and Neck Cancer | Salivary Gland (n=104) | HNSC (n=546) | Upper_aerodigestive_tract (n=669), salivary gland (n=78) |
| Kidney Cancer | Kidney (n=50) | KIRC (n=603) | Kidney (n=1394) |
| Leukemia | Blood (n=2412) | LAML (n=145) | |
| Liver Cancer | Liver (n=188) | LIHC (n=423) | Liver (n=2079) |
| Lung Cancer | Lung (n=607) | LUAD (n=573) | Lung (n=1675) |
| Lymphoma | | DLBC (n=15) | |
| Myeloma | | | |
| Neuroblastoma | Adrenal Gland (n=205) | | Autonomic ganglia (n=393) |
| Ovarian Cancer | Ovary (n=138) | OV (n=265) | Ovary (n=701) |
| Pancreatic Cancer | Pancreas (n=268) | PAAD (n=182) | Pancreas (n=1354) |
| Sarcoma | | SARC (n=265) | |
| Skin Cancer | Skin (n=1000) | SKCM (n=367) | Skin (n=1017) |

**Supplementary Table 4: Mapping of cancer types between different resources**
